# Serum-free differentiation platform for the generation of B lymphocytes and natural killer cells from human CD34+ cord blood progenitors

**DOI:** 10.1101/2025.05.22.655473

**Authors:** Rigveda Bhave, Carla-Johanna Kath, Nadine Rüchel, Eleni Vasileiou, Vera H. Jepsen, Katharina Raba, Aleksandra A. Pandyra, Ute Fischer, Gesine Kogler

## Abstract

**Introduction:** Pre-clinical research on B and NK cell development relies on traditional murine stromal cell-based systems with reduced physiological relevance and clinical applicability.

**Methods:** A serum-free, fully humanized co-culture system utilizing human bone marrow-derived mesenchymal stromal cells (BM-MSCs) was developed to differentiate CB-CD34+ cells towards B and NK cell lineages. Differentiation dynamics were monitored via flow cytometry, with immunophenotypic analysis tracking progression from progenitors to mature cells.

**Results:** The system generated CD19+IgM+ immature B cells and CD56+CD16+ NK cells, recapitulating fetal stages of human lymphopoiesis. Serum-free media conditions ensured reproducibility and high overall yield of B and NK cell progenitors. Flow cytometry identified distinct population peaks, confirming temporal control over differentiation.

**Conclusion:** This clinically relevant platform addresses the limitations of traditional models by providing a more physiologically accurate human microenvironment. The serum-free system supports applications in disease modeling, genotoxic compound screening, and mutational studies of hematopoiesis. By enabling scalable production of B and NK cells it aims to accelerate translational research for immunodeficiencies, cancer immunotherapy, and hematopoietic disorders.

**Graphical Abstract:** 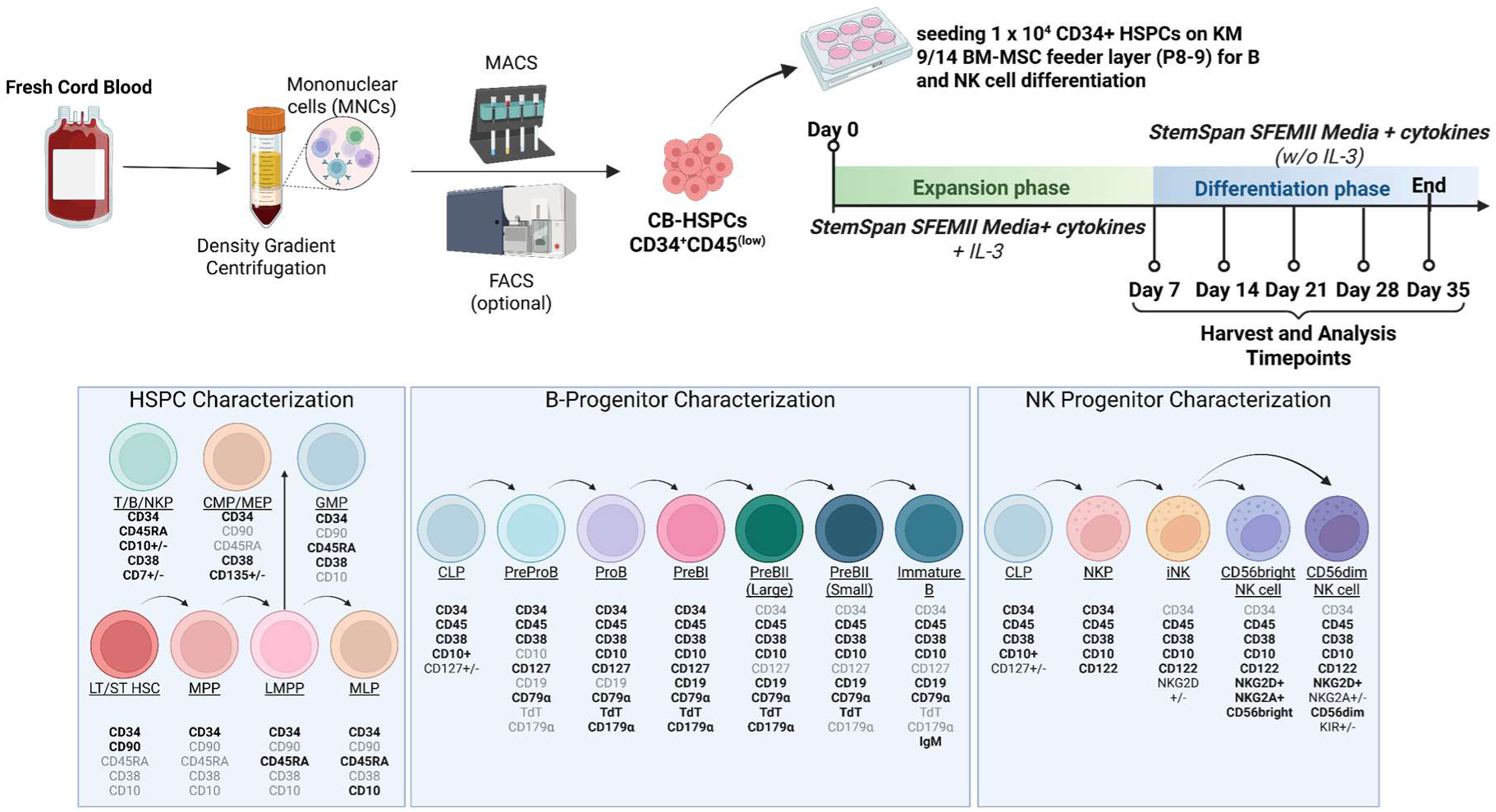

**Significance Statement:** This article presents a novel, fully humanized, serum-free co-culture system that efficiently directs cord blood-derived hematopoietic stem cells into B and natural killer (NK) cells. By using human bone marrow stromal cells and recombinant human cytokines, it overcomes the limitations of murine-based models and better mimics human blood cell development. This platform enables improved disease modeling and therapeutic testing relevant to human hematopoiesis.

## Introduction

Hematopoietic stem and progenitor cells (HSPCs) derived from umbilical cord blood (CB) harbour the potential to self-renew and give rise to all mature blood lineages^1–3^. CB HSPCs are an established source of transplant material for the treatment of hematologic malignancies, bone marrow failure syndromes, hemoglobinopathies and immune deficiencies, where they restore the entire hematopoietic hierarchy in the recipient^4–6^. Several therapies are under development using cells differentiated from CB HSPCs including native/chimeric antigen receptor (CAR) natural killer (NK) cells and CAR T cell therapies^7–9^. Despite these advances, basic translational research frequently utilizes traditional cell culture systems using immortalized murine stromal cells as a supportive microenvironment^10^. These are not conducive to therapeutic/clinical downstream applications and have lower physiological relevance. Moreover, the investigation of B cell and NK cell development is limited by the absence of model systems capable of accurately recapitulating human fetal hematopoiesis under laboratory conditions.

Because B cell precursors of fetal or adult origin are difficult to source in sufficient quantities, the demand for *in vitro* model systems for efficient expansion and differentiation is high. From the earliest CD34+ hematopoietic progenitors to immunophenotypically characterized lineage-restricted cells, the developmental potential towards the B lineage commitment has frequently been evaluated in co-culture systems using the MS-5 murine stromal cell line^11–14^. Previous studies have highlighted the drawbacks of utilizing stromal cells of murine origin, showing that human bone marrow-derived mesenchymal stromal cells (BM-MSCs) have a greater capacity to promote B cell lymphopoiesis^15–17^. While there are multiple high-quality protocols for generating NK cells for clinical applications, these require complex media preparations and often utilize proprietary feeder cell lines and/or specialized coatings^18–20^.

This article presents a novel, user-friendly, serum-free co-culture system for the differentiation of CB-derived CD34+ HSPCs into either CD19+IgM+ immature B cells or CD56+ NK cells, contingent upon the cytokine cocktail employed. The differentiation process is monitored by flow cytometry, enabling precise identification of the time points at which distinct cell populations peak. Immunophenotypic analysis using progenitor-specific markers confirms that this model recapitulates B and NK cell lineage hematopoiesis up to the IgM+ immature B cell and CD56+CD16+ NK cell stages, respectively. In contrast to previously established laboratory-scale models, this approach utilizes a fully humanized system, co-culturing CD34+CD45low HSPCs isolated from cord blood with a human bone marrow-derived mesenchymal stromal cell (BM-MSC) line^21^ and recombinant human cytokines. The serum-free conditions support robust generation of B and NK cell lineage progenitors, facilitating applications in disease modeling, therapeutic and genotoxic compound testing, and mutational studies affecting hematopoietic development.

## Methods

### Materials and Consumables

A complete list of materials is provided in the supplement (Supplemental Table 1.)

### Human CB collection

CB units for research were obtained from donations collected by the José Carreras Cord Blood Bank Düsseldorf with the informed consent of the mothers and ethical approval (Ethic commission votes 2975, 5279, 2020-1144, 2020-9221). All CB units had a gestational age ≥36 weeks.

### CD34+ HSPC isolation and cell counting

Pre-enrichment of the mononuclear cell (MNC) fraction was performed by 1.077 g/cm3 BioColl® (lymphocytes, Bio-Sell) density gradient centrifugation. CD34+ HSPCs were isolated from the MNC fraction using the CD34 MicroBead Kit, UltraPure, Human (Miltenyi Biotech) according to the manufacturer’s instructions. Purity of the CD34+ HSPCs was determined by CD34/CD45/7AAD flow cytometry using a CytoFLEX S Flow Cytometer (Beckman Coulter) according to ISHAGE standards^22^. CD34+ Purity of >80% was considered sufficient for use in experiments, and calculations for cell seeding were adjusted for CD34+ purity.

### Cell counting and proliferation assessment

Total nucleated cell counts (TNCs) of CB and MNCs were determined using an automated hematology analyzer (Cell-Dyn Ruby, Abbott Diagnostics). Manual cell counting of CD34+ HSPCs and differentiated cells in suspension was performed using Trypan Blue dye (Sigma-Aldrich) in a 1:1 ratio with an improved Neubauer chamber slide (NanoEnTek). Cumulative population doublings (CPD) were calculated using the formula: PD = [log(n_­_/n_0_)]/log2; CPD = ∑PD; n_1_ = number of cells counted, n_0_ = number of plated cells.

### BM-MSC feeder cell culture

The human BM-MSC line KM-9/14, derived in-house from a healthy adult donor as previously described^23^, was used at passages P7-P9. Cells were thawed 48 hours before each experiment and plated at 2 x 10^5^ cells per well in 6-well tissue-culture treated plates in Dulbecco’s modified Eagle medium, 1 g/L glucose (DMEM, Gibco) supplemented with 30% fetal bovine serum (FBS, Gibco) and 1% penicillin/streptomycin (Lonza) to achieve 70% confluency on day 0.

### In vitro differentiation cultures

Freshly isolated CB CD34+ HSPCs (1 x 10^4^ to 1 x 10^5^ cells per well) were co-cultured on BM-MSC feeder layers at 70% confluency in 2 mL StemSpan™ SFEM II media (STEMCELL Technologies) supplemented with recombinant human cytokines on day 0. Cytokine cocktails for B and NK cell differentiation are listed in Table 1. On day 7, 2 mL additional media was added, and half-media changes were carried out hereafter every 3-4 days until day 35. To prevent excessive cell death due to HSPC expansion, suspension cell concentrations were maintained below 2 x 10^6^/mL by replating onto fresh feeder cells as required.

**Table 1.** Cytokine cocktails for B and NK cell differentiation.

| HUMAN RECOMBINANT CYTOKINE | FINAL CONCENTRATION |
| --- | --- |
| **B cell differentiation** |  |
| Stem cell factor (SCF) | 50 ng/mL |
| Interleukin-7 (IL-7) | 25 ng/mL |
| Fms-related tyrosine kinase 3 ligand (FLT3-L) | 50 ng/mL |
| Interleukin-3 (IL-3, removed from day 7 onwards) | 10 ng/mL |
| **NK cell differentiation** |  |
| SCF | 20 ng/mL |
| IL-7 | 20 ng/mL |
| FLT3-L | 10 ng/mL |
| IL-3 (removed from day 7 onwards) | 10 ng/mL |
| Interleukin-15 (IL-15) | 10 ng/mL |

### Flow cytometric characterization

To characterize the CB CD34+ HSPCs (Supplemental Figure 1) from various time points during differentiation, the following cell surface marker combinations were used^24^:

hematopoietic Stem Cells (HSCs): LIN-CD34+CD38-CD90+CD45RA-
multipotent progenitors (MPP): LIN-CD34+CD38-CD90-CD45RA-
lympho-myeloid primed progenitors (LMPP): LIN-CD34+CD38-CD90-CD45RA+CD10-
multi-lymphoid progenitors (MLP): LIN-CD34+CD38-CD90-CD45RA+CD10+
pre-B lymphocyte/ NK progenitors (pre-B/NK): LIN-CD34+CD38+CD45RA+/-CD10+
common myeloid progenitors (CMP): LIN-CD34+CD38+CD10-CD45RA-CD135+
megakaryocyte-erythroid progenitors: LIN-CD34+CD38+CD10-CD45RA-CD135-
granulocyte-monocyte progenitors (GMP): LIN-CD34+CD38+CD10-CD45RA+CD135+

Population percentages are expressed as a fraction of the CD34+ live cells, as the BM-MSC feeder cells do not express this marker.

For characterization of the B lineage progenitors arising *in vitro* during differentiation (Supplemental Figure 2), the following surface marker and intracellular marker combinations were applied^25^.

Common lymphoid progenitors (CLP): CD45+CD34+CD38+CD10+CD127+/-CD19-cyCD79α- cyTDT-cyCD179α-

PreProB: CD45+CD34+CD38+CD10-CD127+CD19-cyCD79α+cyTDT-cyCD179α-
ProB: CD45+CD34+CD38+CD10+CD127+CD19-cyCD79α+cyTDT+cyCD179α-
PreBI: CD45+CD34+CD38+CD10+CD127+CD19+cyCD79α+cyTDT+cyCD179α+
PreBII (Large): CD45+CD34-CD38+CD10+CD127-CD19+cyCD79α+cyTDT+cyCD179α+
PreBII (Small): CD45+CD34-CD38+CD10+CD127-CD19+cyCD79α+cyTDT+cyCD179α-
Immature B: CD45+CD34-CD38+CD10+CD127-CD19+cyCD79α+cyTDT-cyCD179α-IgM+

Cytoplasmic antibody staining was performed with the BD IntraSure™ kit (BD Biosciences) as per the manufacturer’s protocol. Population percentages are given as a fraction of the CD38+CD10+ cells, unless stated otherwise.

For characterization of the NK cell progenitors arising *in vitro* during differentiation (Supplemental Figure 3), the following surface marker combinations were used:

NK progenitors (NKP): CD34+CD38+CD45+CD10+CD122+
immature NK cells (iNK): CD34-CD38+CD45+CD10-CD122+NKG2D+/-
CD56Bright NK cell:CD34-CD38+CD45+CD10-CD122+CD56brightCD16+/-NKG2D+NKG2A+KIR-
CD56dim NK cells: CD34-CD38+CD45+CD10-CD122+CD56dimCD16+NKG2D+NKG2A+/-KIR+/-

Population percentages are given as a fraction of the CD45+ live single cells, unless stated otherwise.

Dead cells were excluded using 7AAD (Beckman Coulter) or FVS660/FVS780 (BD Biosciences) depending on the panel (see Supplementary material). Data was acquired on a CytoFLEX S instrument and analyzed using FlowJo™ v10.8 Software (FlowJo LLC). All antibody stainings were performed according to the manufacturer’s protocols in FACS buffer (DPBS+ 2% FBS) at 1:100 (surface staining) and 1:50 (cytoplasmic staining) dilutions. Gating was determined using fluorescence minus one (FMO) and unstained controls at each time point and a combination of compensation beads (BD Biosciences) and CB MNCs were used to establish single-stained controls.

### Flow cytometry-based NK cell cytotoxicity and degranulation assays

CD56+ NK cells were isolated by flow cytometric sorting on day 35 of differentiation and incubated overnight under standard cell culture conditions (37 °C, 5 % CO_2_, humidified atmosphere) in 96-well U-bottom plates. Each well contained 100 µL of NK cell activation medium, consisting of RPMI GlutaMAX supplemented with 10% human AB serum, 30,000 U/mL interleukin-2 (IL-2), and 0.3 ng/mL interleukin-15 (IL-15). NK cell numbers seeded for downstream assays varied depending on the yield obtained from cell sorting. For CB1, 75,000 NK cells were seeded per well (n=3); for CB2, 60,000 NK cells were seeded per well (n=2); and for CB3, 50,000 NK cells were seeded per well (n=2). Monocultures at the corresponding cell densities were included as controls for each CB. On the following day, K-562 target cells (DSMZ, ACC 10) were labeled with 0.17 µM CellTrace™ carboxyfluorescein succinimidyl ester (CFSE, Invitrogen) dye and the staining reaction was quenched by addition of RPMI GlutaMAX supplemented with 10% FBS according to the manufacturer’s instructions. CFSE-labeled K-562 cells were added to the NK cells at an effector-to-target (E: T) ratio of 1:1. Cells were co-cultured for 4 hours at 37 °C and 5% CO₂. Following incubation, supernatants were collected and stored at -80 °C until further analysis by enzyme-linked immunosorbent assay (ELISA). The cells in the pellet were stained for flow cytometric analysis (1:300 antibody dilution, Supplementary Table 1). Target cell killing was quantified as follows: Excess target cell death = % of dead target cells in co-culture - % of dead target cells in monoculture. Degranulation of NK cells during co-culture was measured by cell surface expression of CD107a^26^. NK cell degranulation was calculated as follows: Excess degranulation of NK cells = % of CD107a+ NK cells in co-culture- % of CD107a+ NK cells in monoculture.

### Enzyme-linked immunosorbent assay (ELISA)

Secretion of interferon-gamma (INF-γ) as well as Granzyme B by NK cells was analyzed using the human INF-γ and Granzyme B DuoSet ELISA kit (R&D Systems) following the manufacturer’s instructions. For granzyme B detection, supernatants harvested during NK cell cytotoxicity assays were diluted 1:2. Relative analyte secretion was determined as follows: Relative analyte secretion = C_analyte_ in co-culture/C_analyte_ in monoculture.

### Statistical analysis and figure creation

Row means standard deviation and two-way ANOVA with Dunnet’s multiple comparison tests for flow cytometric data, and statistical analysis of NK cell cytotoxicity assays and ELISAs were performed using GraphPad Prism v8.0.2 (GraphPad Software). Image creation and figure assembly was performed on Inkscape v1.4 (The Inkscape Project). The graphical abstract was created using BioRender (https://www.biorender.com/).

## Results

### CB CD34+ HSPCs expand in B differentiation conditions and generate lymphoid-primed progenitors

There is substantial variability in the percentage of CD34+ cells among different CB units, influenced by factors such as gestational age, birth conditions and inherent biological heterogeneity^27–29^. CD34+CD45low cells were isolated from n=9 biologically distinct CB units. Given the importance of generating sufficient differentiated hematopoietic cells from a limited CD34+ cell input, hematopoietic cell proliferation was monitored throughout the B cell differentiation culture period. Since hematopoietic cells only loosely adhere to the BM-MSC feeder layer, manual cell counts were conducted by gently resuspending the cells in the well without disturbing the feeder layer. During later time points, detachment of non-viable feeder cells was observed; however, these cells could be excluded from the count based on their larger size and positive Trypan blue staining. By day 35, a cumulative population doubling of 44.69±0.90 was observed in hematopoietic cell numbers, indicating robust generation of progenitors (Figure 1A).

**Figure 1.**
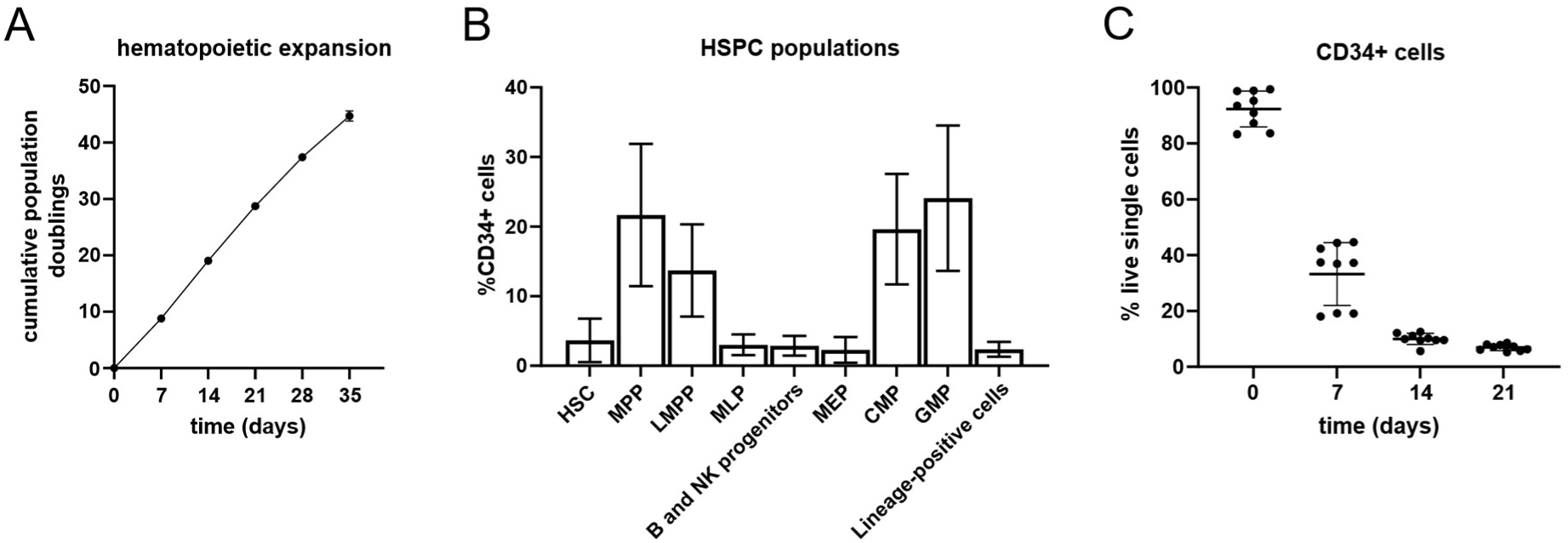
CB CD34+ HSPC dynamics in B cell differentiation culture conditions. (A) Expansion of hematopoietic cells during B cell differentiation depicted as the cumulative population doubling over time. (B) Immunophenotypic characterization of CD34+ progenitor populations in freshly isolated CB HSPCs on day 0. (C) Fraction of CD34+ cells measured at experimental time points as a percentage of the live cells. Error bars indicate mean ±SD of n=9 independent biological CB units.

There were significant differences between the yields and composition of CD34+CD45low HSPCs isolated on day 0 (Figure 1B). Immunophenotypic characterization of the HSPC progenitor populations on day 0 showed that the majority of CD34+ cells isolated from CB were MPPs (CD34+CD38-CD90-) (21.68±10.23%), CMPs (CD34+CD38+CD135+) (19.62±7.92%) and GMPs (CD34+CD38+CD135+CD45RA+) (24.08±10.44%) (Figure 1B). Overall, CB samples exhibited a myeloid progenitor (CMP, GMP, MEP) composition between 45-60%. After day 21, the fraction of CD34+ cells that were not already lineage committed (CD34+LIN+) was negligible, and therefore not analyzed (Figure 1C).

Under B cell differentiation conditions, the HSCs (CD34+CD38-CD90+), MPPs and LMPPs (CD34+CD38-CD90- CD45RA+) (Figure 2A, first row, 2B) generated CD38- early lymphoid-primed MLP(CD34+CD38-CD45RA+CD10+) population peaks on day 7 (22.98±17.02%) and day 14 (20.78±9.03%). By day 21, the early hematopoietic progenitors were pushed to proliferate towards the CD10+ lymphoid primed B/NK progenitor state (74.99±7.65%) (Figure 2A, third row, 2C), accompanied by a significant reduction in the MEP (CD34+CD38+CD135-CD45RA-), CMP and GMP myeloid progenitor populations (Figure 2A, fourth row, 2D).

**Figure 2.**
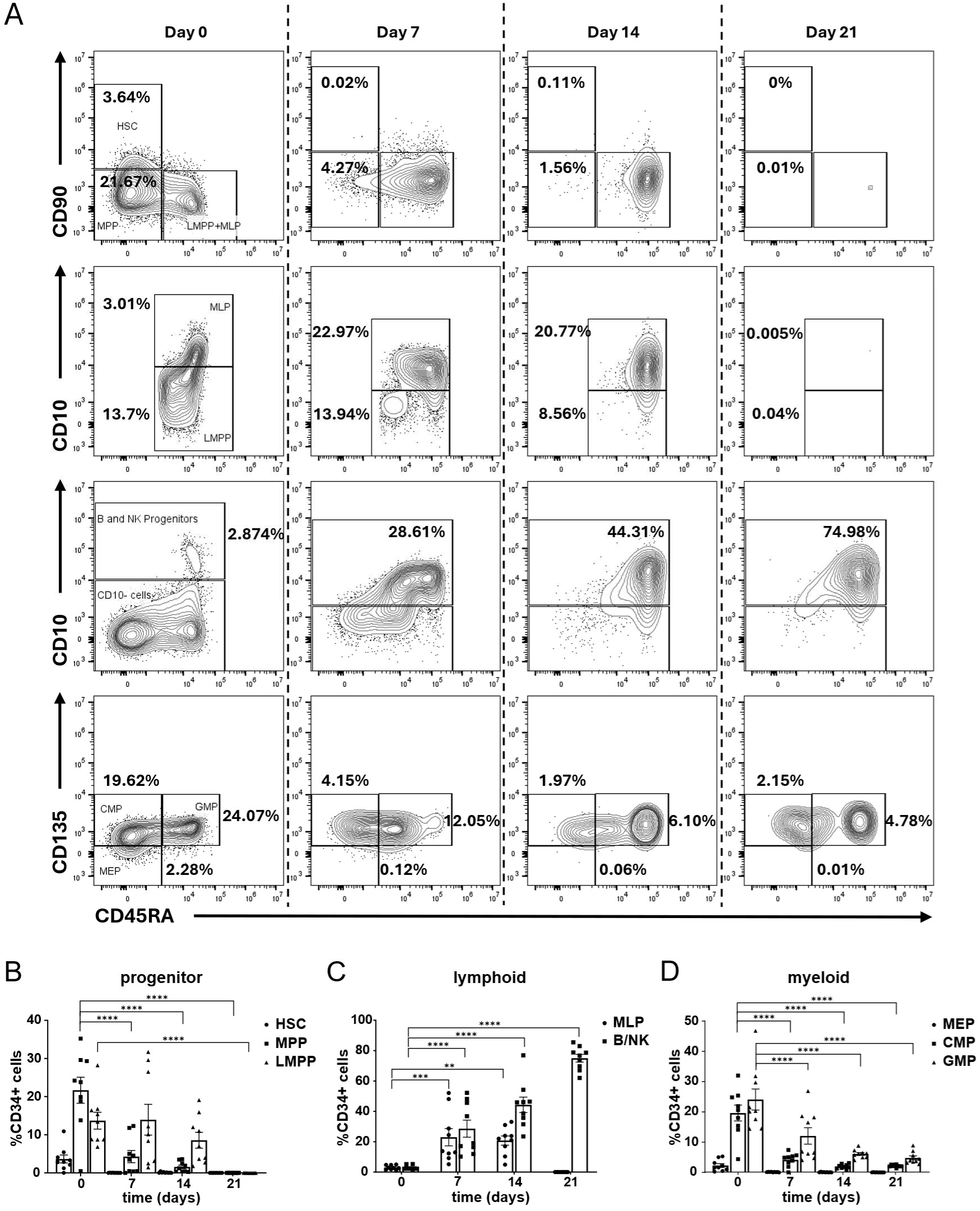
Development of LIN-CD34+ hematopoietic progenitor cells in B cell culture conditions. (A) First row: CD90 expression discriminates between the earliest HSC population and the MPPs in the CD34+CD38- compartment. Second row: CD10 expression distinguishes the CD45RA+ cells into LMPPs and MLPs in the CD34+CD38- compartment. Third row: CD10 expression separates the CD34+CD38+ compartment into the CD10+ B and NK cell progenitors and myeloid progenitors. Fourth Row: The CD10- compartment consists of the MEPs, CMPs and GMPs. Percentages indicated as a proportion of CD34+ cells at each experimental time point. (B) Distribution of the earliest hematopoietic progenitors (HSCs, MPPs and LMPPs), (C) lymphoid-primed progenitors expressing CD10 (MLPs, B/NK progenitors), and (D) myeloid-primed progenitors. Error bars indicate the mean±SD of n=9 biological replicates. A two-way ANOVA for multiple comparisons was performed to determine statistical significance (**p<0.01, ***p<0.001, ****p<0.0001).

### B cell differentiation generates CD38+CD10+ early and CD19+ late committed progenitors *in vitro*

The CD38+CD10+ CLP state marks the beginning of lymphoid commitment in hematopoiesis^30^. During *in vitro* B cell differentiation, the CD38+CD10+ population was already established (56.14±15.06%) by day 7 (Figure 3A, first row, 3B), the majority of which were CD34+ CLPs (29.59±18.20%) (Figure 3C). The CD10-PreProB (1.24±0.72% of CD45+ live cells on day 21) and CD10+ ProB (1.06±0.006%) populations, identified as the earliest B-committed fetal progenitors^31^, were also detectable (Figure 3A, C, D). They are distinguished from the CLPs by expression of the early pre-B cell receptor (pre-BCR) subunit CD79α. The pre-BCR complex is essential for quality control during differentiation, allowing only cells with a successfully rearranged µH chain to progress further^32^. Notably, the PreProB population was CD45low, similar to early CD34+ hematopoietic progenitors. The remaining CD38+CD10- cells were identified as CD33+CD11b+ expressing monocytes, which were observed to form aggregates in culture if the cell density was too high (data not shown).

**Figure 3.**
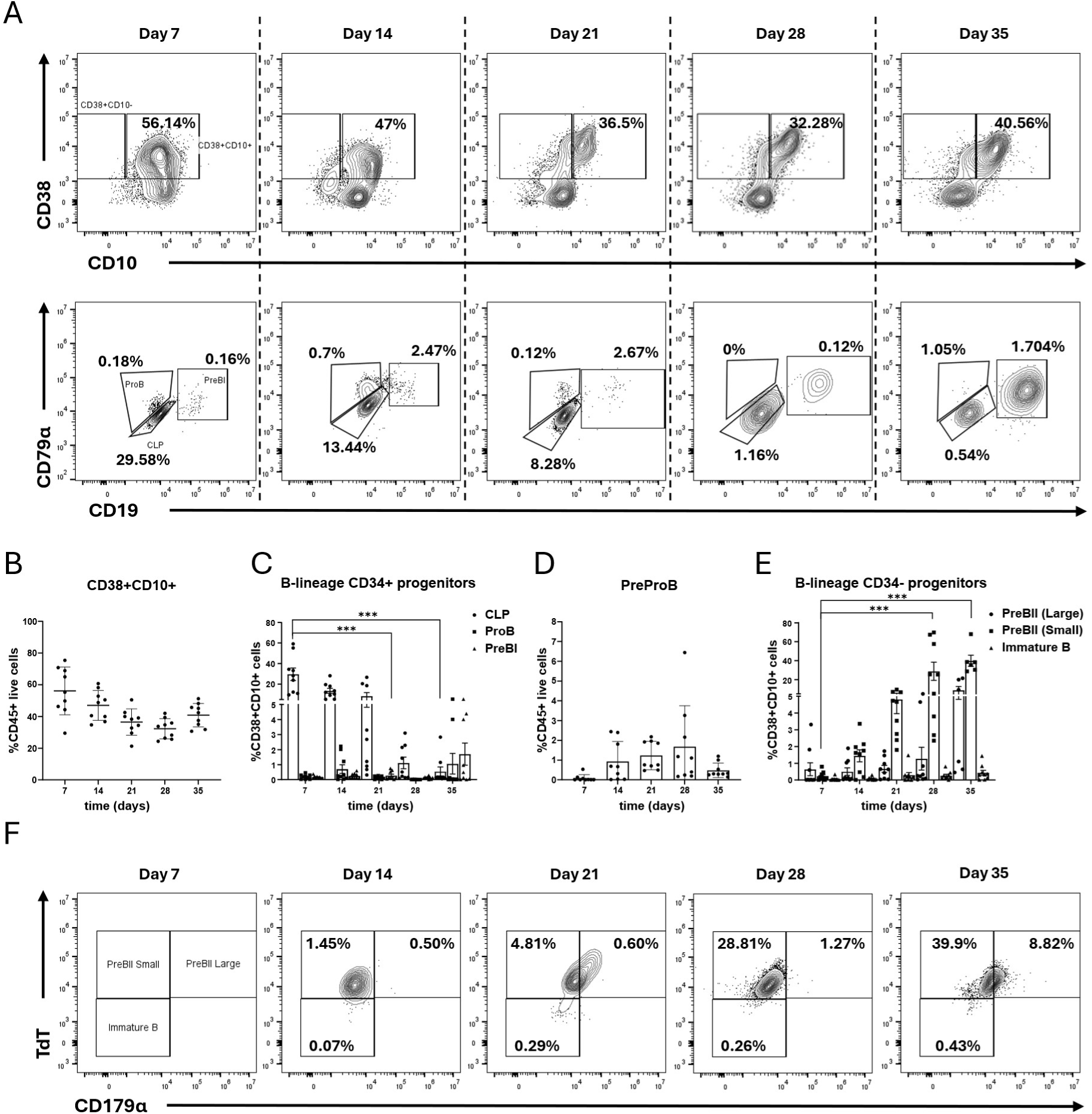
Early B lineage committed progenitors arising in co-culture conditions. (A) First row: CD38+CD10+ cells include all lineage-committed B progenitors from the CLP to the IgM-expressing immature B cell. Percentages shown are as a proportion of CD45+ live single cells at each experimental time point. Second row: The early B progenitors still express CD34. The CLPs give rise to the ProB cells, which express CD79α and CD179α, and upon gaining CD19 expression become PreBI cells. Percentages shown are as a proportion of CD38+CD10+ cells at each experimental time point. (B) CD38+CD10+ cells as a percentage of the CD45+ hematopoietic cells in culture. (C) Emergence of CD34+ B progenitors as a percentage of CD38+CD10+ cells. (D) PreProB is the earliest known B-committed progenitor, and resides in the CD38+CD10- compartment, here shown as a percentage of the CD45+ live hematopoietic cells. (E) The CD34-CD19+ compartment consists of the PreBII small/large and the Immature B cells. Error bars indicate the mean±SD of n=9 biological replicates. A two-way ANOVA for multiple comparisons was performed to determine statistical significance (***p<0.001). (F) Emergence of CD34-CD19+ B populations (PreBII(large), PreBII(small) and ImmatureB) over 35 days. Percentages shown are as a proportion of CD38+CD10+ cells at each experimental time point.

The earliest B progenitor to express CD19 was identified as the PreBI cell, which arose in small numbers throughout the differentiation process and peaked on day 35 (1.70±2.19%) (Figure 3A, C). The CD34-CD19+ PreBII (large) and PreBII (small) cells were distinguished by the expression of CD179α, and these two populations represented the majority of CD19+ cells by day 35 (Figure 3A, second row, 3E, F). Only a small fraction of cells (0.23±0.05%) expressing IgM were observed at on day 35, although it was not attempted to extend the cultures beyond this timepoint. Hence it was established that the B cell differentiation model system could successfully drive CB CD34+ HSPCs towards CD19+ expressing B cells, and the progenitor populations could be immunophenotypically identified.

### IL-15 supplementation drives CB-derived CD34+ HSPCs to differentiate towards NK cells

IL-15 promotes NK cell development by activating key signaling pathways including JAK-STAT5, PI3K-AKT-mTOR, and RAS-MEK-MAPK, driving NK cell maturation, survival and function^33, 34^. Using a modified cytokine cocktail for NK cell specific differentiation (Table 1), significant quantities of hematopoietic cells were generated in the NK cell differentiation system (CPD=10.66±0.57) (Figure 4A) across n=6 independent biological CB replicates. Compared to the B cell differentiation system, the population doublings were lower, however this was attributed to the lower concentrations of SCF and FLT3-L (Table 1), which promote expansion of early hematopoietic progenitors.

**Figure 4.**
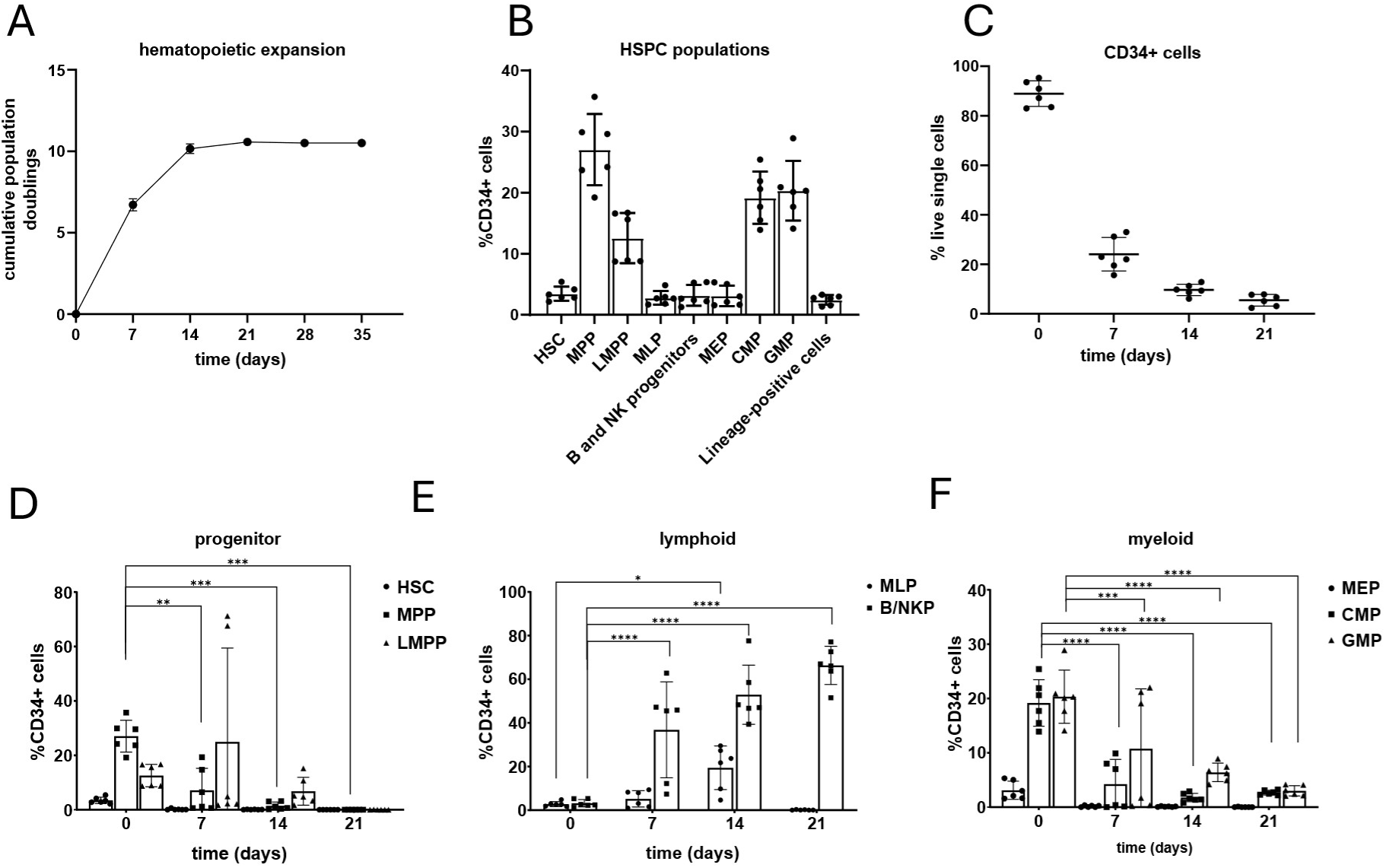
Development of CD34+ hematopoietic progenitor cells in NK cell culture conditions. (A) Expansion of hematopoietic cells during NK cell differentiation depicted as the cumulative population doubling over time. (B) Immunophenotypic characterization of CD34+ progenitor populations found in freshly isolated CB HSPCs. (C) Fraction of CD34+ cells measured at experimental time points as a percentage of the live cells. (D) Distribution of the earliest hematopoietic progenitors (HSCs, MPPs and LMPPs), (E) lymphoid-primed progenitors expressing CD10 (MLPa, B/NK progenitors), and (F) myeloid-primed progenitors (MEPs, CMPs and GMPs). Error bars indicate the mean±SD of n=6 biological replicates. A two-way ANOVA for multiple comparisons was performed to determine statistical significance (*p<0.05, **p<0.01, ***p<0.001, ****p<0.0001).

As previously mentioned, heterogeneity was found between the CB samples upon immunophenotyping of the HSPC lineages (gated as in Figure 2A, Supplemental Figure 1) on day 0, with enrichment in the MPP, CMP and GMP fractions (Figure 4B). Under NK cell differentiation conditions, there was a significant reduction in CD34+ progenitors from day 0 (88.91±5.1%) to day 21 (5.44±2.35%) (Figure 4C). The early hematopoietic progenitors (HSCs, MPPs, LMPPs) were lost by day 14 (Figure 4D). This exhaustion of the early hematopoietic progenitors from the culture system also seems to be a contributing factor to limited cell expansion compared to the B cell differentiation system. There was a skew towards the CD10+ B/NKP cells by 21 days (66.33±8.74%) (Figure 4E), and a significant reduction in CMP (19.16±4.28% to 2.99±0.43%) and GMP (20.33±4.89% to 3.00±0.95%) myeloid progenitors (Figure 4F). It was established that the addition of IL-15 to the cytokine cocktail had no detrimental impact on the HSPC populations heading towards the lymphoid lineage.

The key stages of NK cell differentiation from HSCs through the CLP, NKP and iNK stages have been previously described^35^. A flow cytometric strategy (Figure 5A, B, Supplementary Figure 3) to identify these populations in the NK cell co-culture system was employed. CLPs were detectable on day 7 (5.27±1.03%) (Figure 5B, E). NKPs, which are characterized by expression of CD122, were also most abundant at day 7 (3.28±1.11%) (Figure 5B, E). There were most likely higher percentages of these progenitors prior to day 7, but due to the significant loss of CD34+ populations after day 7 CLPs and NKPs were found only in low numbers after this time point. The iNK cells, found in the fetal liver during early development^36^, were detected in significant numbers on day 7 (24.94±6.14%), and day 14 (27.15±9.41%) (Figure 5C, E). Definitive CD56+ NK cells began to emerge as early as day 14 (0.81±0.76%) in culture and reached a peak at day 35 (28.46±7.01%) (Figure 5 A, E). It was not possible to discern between the dominant CD56bright and the more mature but rarer CD56dim populations due to panel limitations, but analysis of CD16 expression demonstrated the emergence of a more cytotoxic CD56+CD16+ NK cell population by day 35 (30.26±9.22% of CD56+ cells) (Figure 5D, G). Other NK-specific markers, i.e. KIR2DL1, NKG2D and NKG2A, were detectable on the CD56+ cells starting Day 21 (Figure 5F). As with the B cell differentiation, the remaining cells in culture were identified as CD33+CD11b+ expressing monocytes, with similar aggregation characteristics at high cell densities (data not shown).

**Figure 5.**
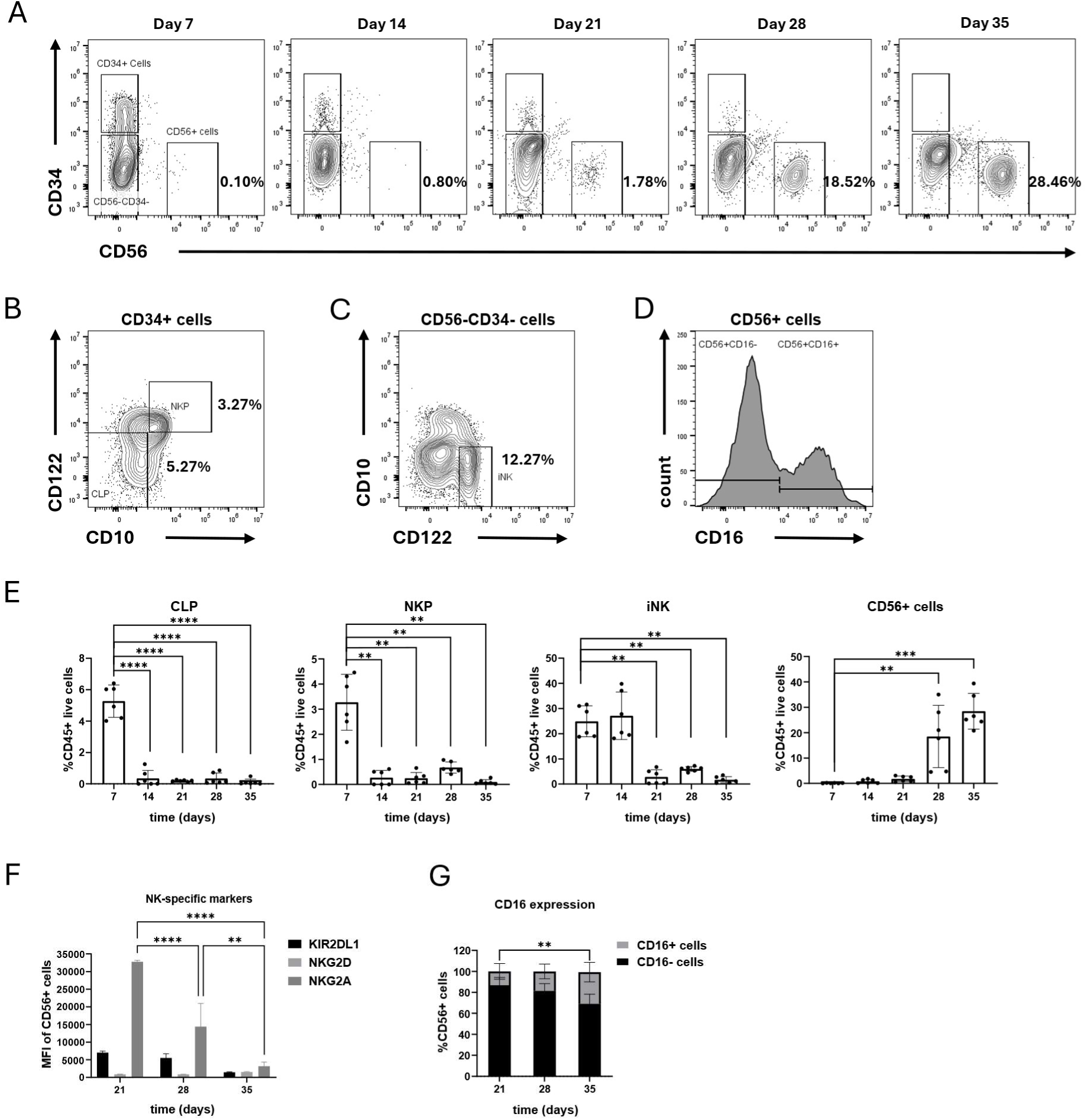
Differentiation of CB HSPCs towards CD56+ NK cells in co-culture system. (A) Representative flow cytometry plots demonstrating initial loss of CD34+ HSPCs and emergence of CD56+ NK cells over 35 days. Plots are gated on the CD45+ live populations, and sub-gated to identify (B) CLP, NKP and (C) iNK progenitor populations. Percentages inside plots indicate the mean proportions as a fraction of CD45+ live cells. (D) Histogram of CD16 expressing CD56+ cells. (E) Emergence of various NK cell populations over the course of 35 days. (F) Mean fluorescence intensities (MFI) of NK-specific markers expressed on CD56+ cells. (G) CD56+ cells begin expressing CD16 from day 21 onwards. Error bars indicate mean±SD of n=6 independent biological CB samples. A two-way ANOVA for multiple comparisons was performed to determine statistical significance (**p<0.01, ***p<0.001, ****p<0.0001).

### NK cells generated *in vitro* effectively kill K-562 target cells in co-culture

The cytotoxic potential of differentiated NK cells was evaluated using a standard killing assay^37^. Differentiated NK cells were first isolated by fluorescence-activated cell sorting (FACS) and subsequently co-cultured with the K-562 target cell line. Following co-culture, both NK and target cells were analysed by flow cytometry to assess target cell death, NK cell degranulation, and the expression of surface markers relevant to NK–target cell interactions. NK cell-mediated cytotoxicity resulted in an excess target cell death rate of 55.37±7.77%, while NK cell degranulation increased by 14.24±2.80% (Figure 6A). The relative MFI of most surface markers remained largely unchanged, apart from PD-L1 (MFI 5.55±2.47) (Figure 6B). To further quantify cytokine and effector molecule release, supernatants from the co-cultures were subjected to ELISA for IFN-γ and Granzyme B detection. The relative analyte secretion levels of IFN-γ (2.41±1.47) and Granzyme B (1.58±0.22) were found to be modestly elevated (Figure 6C). These results show that NK cells produced in this co-culture system have active cytotoxic activity and can secrete cytokines upon stimulation.

**Figure 6.**
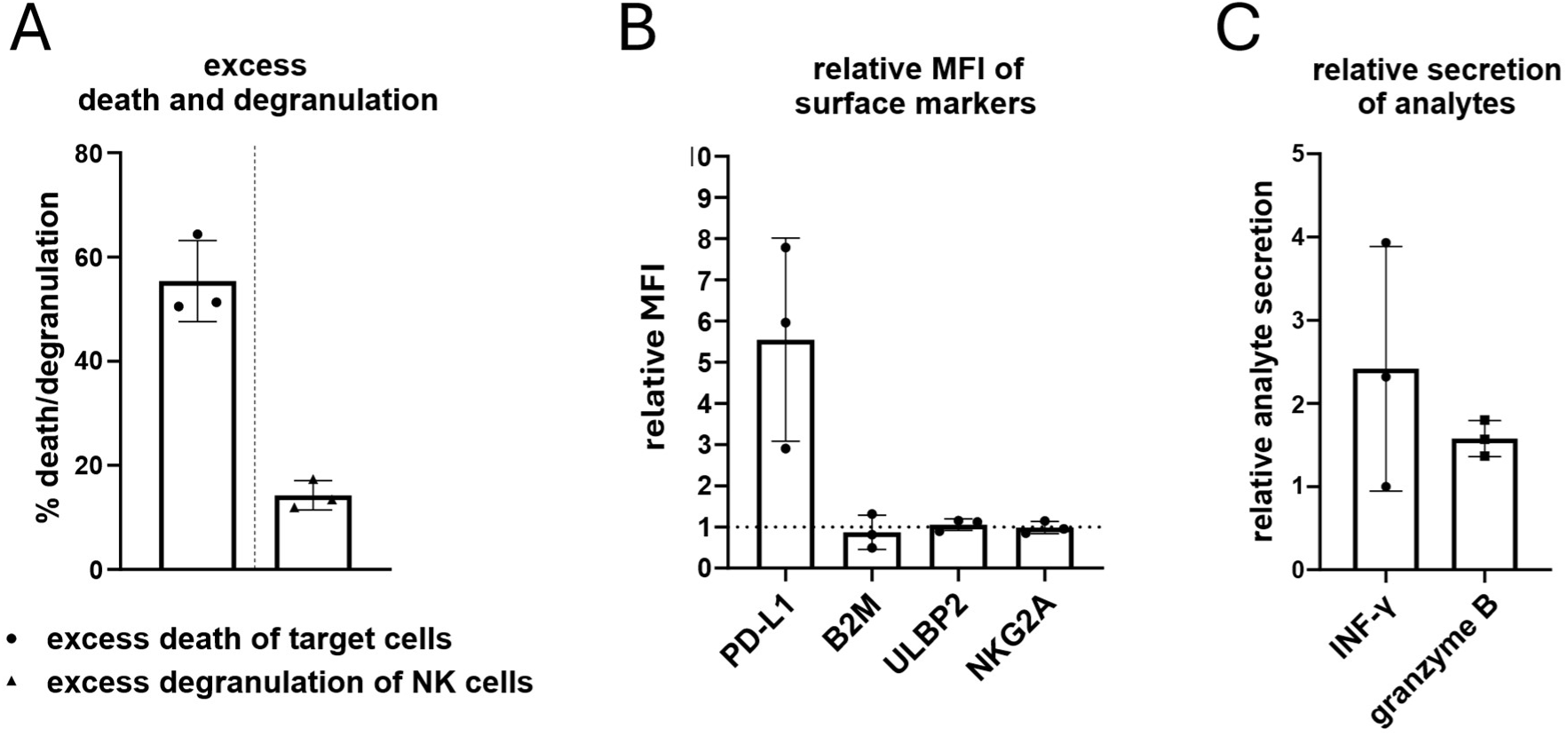
Cytotoxic potential of day 35 CD56+ sorted NK cells. (A) Percentages of excess K-562 target cell death and NK cell degranulation marked by CD107a expression. (B) Relative MFI of major surface markers involved in NK cell killing activity. (C) Relative secretion of analytes into the supernatant of co-cultured cells. Error bars indicate the mean±SD of n=3 independent biological CB samples.

## Discussion

In response to a necessity for a straightforward yet versatile model system that can recapitulate fetal B cell and NK cell development within pre-clinical research settings, the objective was to establish a protocol to generate B and NK cells in sufficient quantities for experimentation. Additionally, we aimed to develop a robust approach to identify and characterize emerging progenitor populations during *in vitro* culture. Recent single-cell transcriptomic studies demonstrated more complex heterogeneity in human hematopoietic precursors than previously understood, emphasizing the necessity for a more detailed characterization of emerging progenitor populations in *in vitro* differentiation systems^38, 39^. Here, we demonstrate (1) a humanized, serum-free stromal co-culture system enabling scalable production of early/late B cell progenitors from human CB HSPCs, (2) directed differentiation towards CD56+ NK cells via exogenous IL-15 supplementation, and (3) streamlined flow cytometric approaches to track hematopoietic populations developing *in vitro*.

B cell differentiation systems utilizing co-culture with murine stromal cells^40^ have substantially advanced research across multiple disciplines, including developmental biology^41, 42^ and pediatric leukemia^43^. These stromal feeder layers provide a supportive microenvironment that mimics *in vivo* conditions, enabling the proliferation, survival and differentiation of B cell progenitors through both direct cell-cell interactions and the secretion of soluble factors such as cytokines^44, 45^. Current understanding of hematopoietic progenitor populations is primarily derived from murine-based models^46^, despite documented functional disparities between murine and human stromal cells in HSPC maintenance and differentiation^47, 48^. It was previously demonstrated that using human BM-MSCs as feeder can produce B lymphocytes from CB CD34+ cells^49^. Although these studies successfully generated CD19+ cells, in many instances, high purities did not translate into fold-expansion numbers necessary for downstream applications. The serum-free formulations allow for greater control over the differentiation process, as well as reproducibility for downstream applications by minimizing interactions with genotoxic/therapeutic compounds, that may adversely react to the presence of serum^50^. In addition, serum-free differentiation systems are more amenable to scale-up for large-scale cell production and regulatory compliance. Maintaining the system within budgetary constraints and at a practical scale for laboratory use remained key considerations in the development of this differentiation model.

As previously noted in the results, CB CD34+ HSPCs are enriched for MPPs and myeloid progenitors. By following the differentiation process immunophenotypically, it was demonstrated that despite inherent biological variability, the differentiation conditions preferentially promoted the expansion of CD10+ progenitors, subsequently generating CD19+ B cell populations. Analysis of stage-specific marker expression confirmed that all phases of B cell development up to the IgM+ stage were recapitulated in this system. The capacity to repeatedly passage earlier progenitors (day 14 and earlier) onto multiple stromal feeder layers demonstrated the robust scalability of the differentiation protocol. It is important to note that permitting uncontrolled expansion of hematopoietic progenitors within a confined culture environment may result in rapid cell death and impaired B lineage cell production, favouring differentiation towards monocytes instead^51^.

In contrast, NK cells have been efficiently differentiated *in vitro* from CB CD34+ cells, with excellent yields obtained from good manufacturing practice compliant protocols^5, 9, 20^. This is increasingly important for immunotherapy due to the need for large numbers of functional NK cells and the advantages offered by CB as a source^52^. A modification to the B cell differentiation system with the addition of IL-15 over five weeks was made. This approach enabled a reduction in the costs associated with establishing a new model system, while still attaining a high overall fold expansion. Importantly, the functional properties of the NK cells were in line with previously described protocols, demonstrating cytotoxic activity towards K-562 target cells as well as NK cell degranulation^18, 52^. Further optimization of the media conditions, and the addition of cytokines such as IL-2 are intended to increase the CD56+ yield even further and ameliorate the rapid loss of CD34+ progenitors at the earlier time points^53^. The relatively low proportions of NK cell progenitor populations in proportion to CD56+ emergence may be addressed by further immunophenotypic identification, such as CD7+ ProT populations^54^. It must be mentioned here that feeder-free protocols for both B and NK cell differentiation are available, however they require the establishment of two separate culture systems^54, 55^. Future studies aim to validate this model for use with induced pluripotent stem cell-derived HSPCs, to explore developmental susceptibility of progenitor cells to oncogene-induced processes^56^.

In conclusion, the results presented above suggest that the serum-free, co-culture system presented here provides a developmentally relevant setting for the examination of B and NK cell progenitors with a minimal set of requirements. This system requires a low input of CB HSPCs, factors in donor variability and consistently generates robust yields of hematopoietic progenitors that can be characterized by flow cytometry. This system will be used to analyse the lymphoid differentiation characteristics of different populations of progenitors arising from preleukemic translocations in early progenitors^57, 58^, and to identify defects in B and NK cell lymphopoiesis induced by exogenous environmental toxins^59^. It is also intended to further characterize sorted populations at bulk and single-cell RNA transcriptomic level.

## Data Availability

The data underlying this article are available in the article and in the supplementary material.

## Acknowledgements

The authors would like to thank Janine Korn, Almuth Düppers and Dr. Stefanie Liedtke at the Institute for Transplantation Diagnostics and Cell Therapeutics (Heinrich Heine University, Medical Faculty, Düsseldorf, Germany) for their assistance and expertise in the handling and processing of cord bloods. We would also like to thank the José-Carreras Cord Blood Bank, Düsseldorf, for providing infrastructure and access to primary cord blood materials.

