## Supplemental material for "Serum-free differentiation platform for the generation of B lymphocytes and natural killer cells from human CD34+ cord blood progenitors"

#### Supplemental tables

*Supplemental Table 1. List of materials.*

| REAGENT or RESOURCE | SOURCE | IDENTIFIER |
|---|---|---|
| Antibodies (HSPC Panel) | | |
| 7AAD | Beckman Coulter | A07704 |
| CD10-PC7 | BioLegend | 312214 |
| CD127-R718 | BD Biosciences | 566967 |
| CD135-PE | BioLegend | 313306 |
| CD15-BV510 | BioLegend | 323028 |
| CD34-Pacific Blue | BioLegend | 343512 |
| CD38-BV650 | BD Biosciences | 569966 |
| CD45RA-APCFire750 | BioLegend | 304152 |
| CD7-APC | BioLegend | 982702 |
| CD90-FITC | BioLegend | 328108 |
| Lineage Cocktail-BV510 | BioLegend | 348807 |
| Antibodies (B cell Panel) | | |
| CD10-APCFire750 | BioLegend | 312230 |
| CD127-R718 | BD Biosciences | 566967 |
| CD179α-BV421 | BD Biosciences | 566583 |
| CD19-RB705 | BD Biosciences | 570235 |
| CD34-RB780 | BD Biosciences | 569078 |
| CD38-BV650 | BD Biosciences | 569966 |
| CD45-BV786 | BD Biosciences | 563716 |
| CD79a-PE | BD Biosciences | 555935 |
| FVS660 | BD Biosciences | 564405 |
| IgM-BV510 | BD Biosciences | 563113 |
| TdT-FITC | BD Biosciences | 332789 |
| Antibodies (NK Panel) | | |
| CD10-PC7 | BD Biosciences | 565282 |
| CD122-APC | BioLegend | 339008 |

| | | |
|---|---|---|
| CD158a-PE | BD Biosciences | 556063 |
| CD159a-BV786 | BD Biosciences | 747917 |
| CD16-RB705 | BD Biosciences | 570243 |
| CD314 (NKG2D)- BV421 | BioLegend | 320822 |
| CD34-PE/Dazzle594 | BioLegend | 343534 |
| CD38-BV560 | BD Biosciences | 569966 |
| CD3-APCFire750 | BioLegend | 300470 |
| CD45-FITC | Beckman Coulter | 345808 |
| CD56-BV510 | BioLegend | 318340 |
| FVS780 | BD Biosciences | 565388 |
| Antibodies (NK Killing assay) | | |
| PD-L1-PE/Dazzle594 | BioLegend | 329732 |
| B2M-PerCP-Cy5.5 | BD Biosciences | 656645 |
| CD3-PC7 | eBioscience | 25-0038-42 |
| CD56-AF647 | BD Biosciences | 557711 |
| CD107a-APCCy7 | BioLegend | 328630 |
| ULBP2-BV605 | BD Biosciences | 748131 |
| CD314 (NKG2D)-BV786 | BD Biosciences | 743560 |
| Other flow cytometry reagents | | |
| BD Intrasure™ Kit | BD Biosciences | 641778 |
| Brilliant Stain Buffer | BD Biosciences | 563794 |
| Fc Block | BD Biosciences | 564219 |
| VersaLyse lysing solution | Beckman Coulter | A09777 |
| DAPI | Invitrogen | D1306 |
| Critical commercial assays | | |
| CD34 MicroBead Kit UltraPure | Miltenyi Biotec | 130-100-453 |
| CellTrace™ CFSE Cell Proliferation Kit | Invitrogen | C34554 |
| Human Granzyme B DuoSet ELISA | R&D Systems | DY2906 |
| Human IFN-gamma DuoSet ELISA | R&D Systems | DY285B |

| | | |
|---|---|---|
| LS Columns | Miltenyi Biotec | 130-042-401 |
| MACS® MultiStand | Miltenyi Biotec | 130-042-303 |
| Pre-Separation Filters (30 µm) | Miltenyi Biotec | 130-041-407 |
| Chemicals and Recombinant Proteins | | |
| Albunorm® 20% | OctaPharma | - |
| Biocoll® (Lymphocytes) | Bio&Sell | BS.L6115 |
| C-Chip (4ch) Hemocytometer | NanoEntek | DHC-NO4 |
| CliniMACS® PBS/EDTA Buffer | Miltenyi Biotec | 700-25 |
| Dimethyl Sulfoxide (DMSO) | Sigma Aldrich | D2650-100ML |
| DMEM | Gibco | 31885-023 |
| Dulbecco's Phosphate Buffered Saline (PBS), without calcium and magnesium | Sigma-Aldrich | D8537-500ML |
| FCS | Gibco | A5256701 |
| Human Recombinant Flt3/Flk-2 Ligand | STEMCELL Technologies | 78009 |
| Human Recombinant IL-15 | STEMCELL Technologies | 78031 |
| Human Recombinant IL-3 | STEMCELL Technologies | 78040 |
| Human Recombinant IL-7 | STEMCELL Technologies | 78053 |
| Human Recombinant SCF | STEMCELL Technologies | 78062 |
| Pencillin/Streptomycin | Gibco | 15140122 |
| RPMI GlutaMAX | Gibco | 61870036 |
| StemSpan SFEM II | STEMCELL Technologies | 09655 |
| Trypan Blue Stain | Gibco | 15250-061 |
| TrypLE™ Select (1x) | Gibco | 12563-029 |
| Instruments | | |

| | | |
|---|---|---|
| Cell-Dyn Ruby | Abbott Diagnostics | 08H67-01 |
| CytoFlex S (V5-B5-R3) | Beckman Coulter | C09734 |
| Software and Algorithms | | |
| CytExpert v2.6 | Beckman Coulter | https://www.beckman.com/flow-cytometry/research-flow-cytometers/cytoflex/software |
| FlowJo v10.8 | FlowJo LLC | http://www.flowjo.com/ |
| GraphPad Prism v8.0.2 | GraphPad Software | https://www.graphpad.com/ |
| Inkscape v1.4.2 | The Inkscape Project | https://inkscape.org/ |
| BioRender Software | BioRender | https://www.biorender.com/ |

### Supplemental Figures

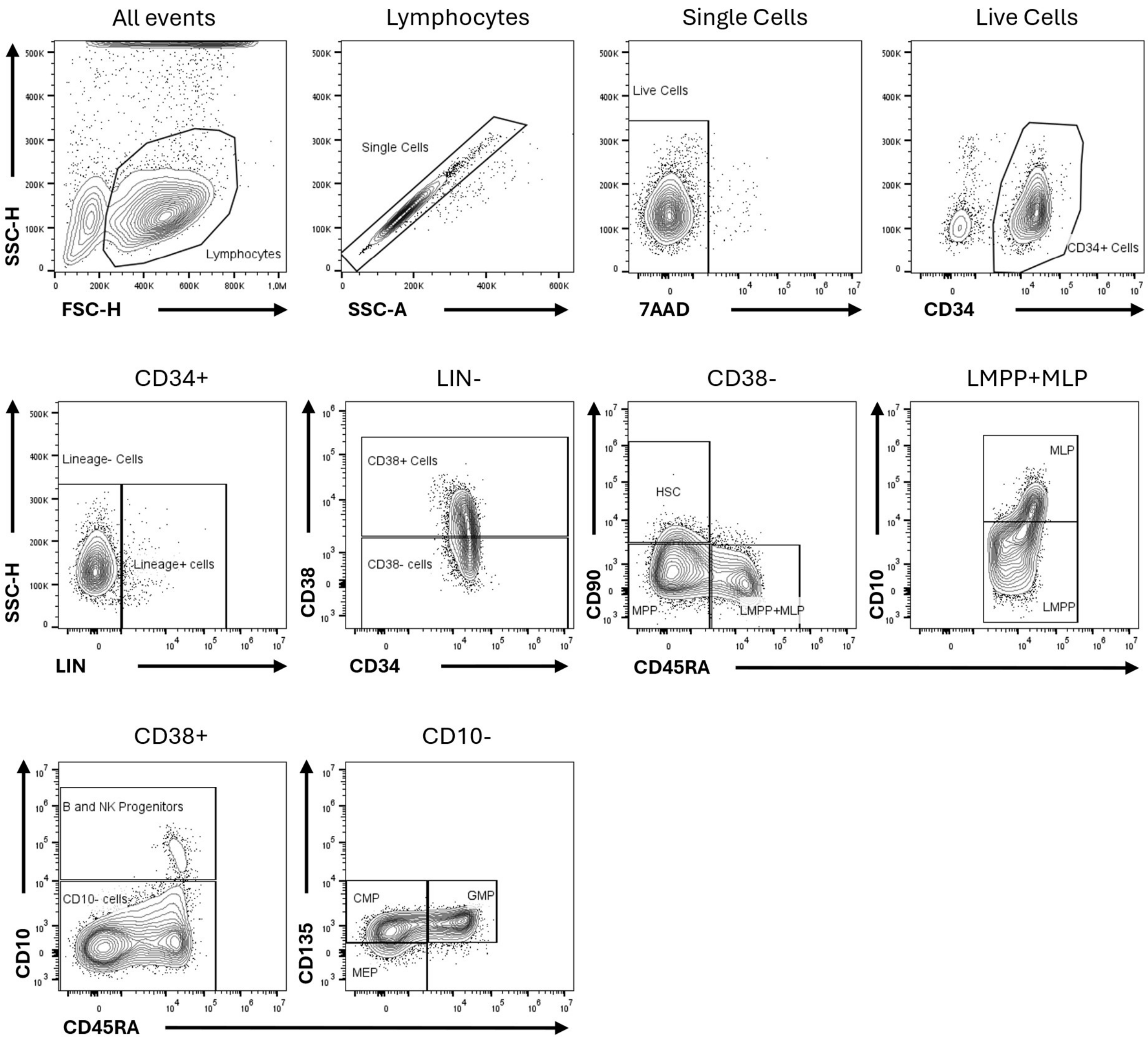


*Supplemental Figure 1. **Flow cytometric analysis of CD34+ HSPC progenitors during B and NK differentiation.** HSPC Panel successive gating strategy used for flow cytometric identification of HSPC progenitors. Plots are representative of freshly isolated day 0 CB CD34+ cells.*

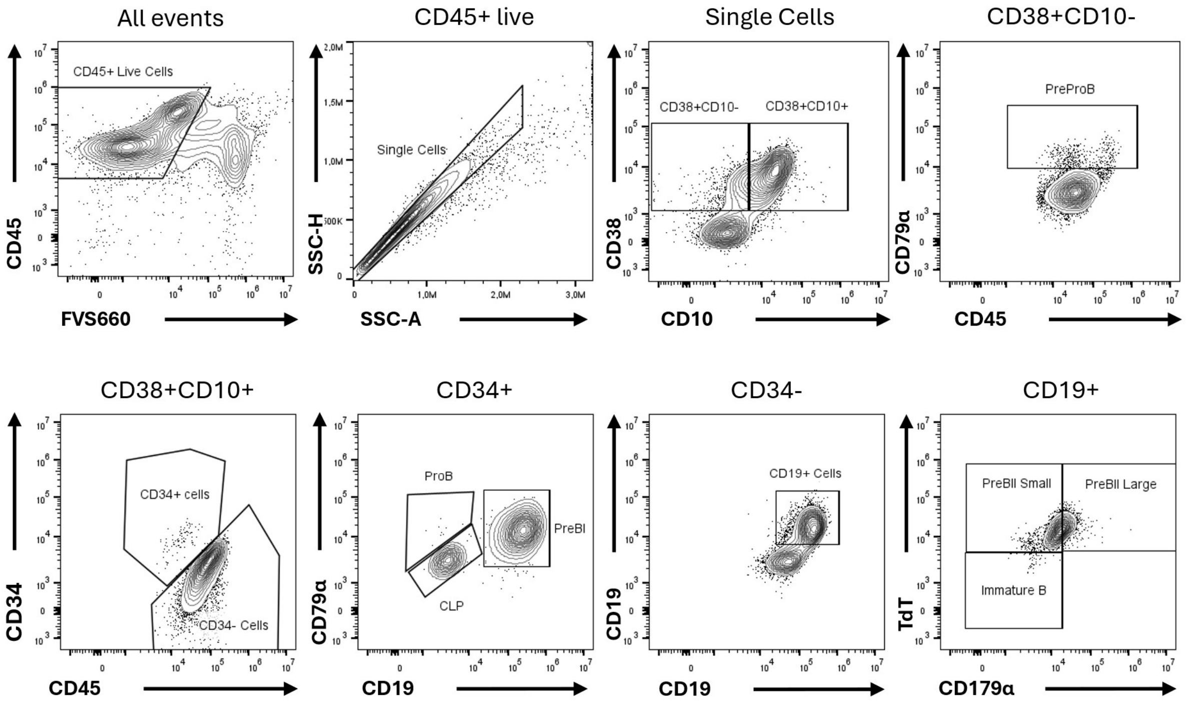


*Supplemental Figure 2. Flow cytometric analysis of B lineage progenitors during B cell* ***differentiation.*** *B cell panel successive gating strategy used for immunophenotypic identification of B lineage progenitors. Plots are representative of cells analyzed on day 35 of B cell differentiation.*

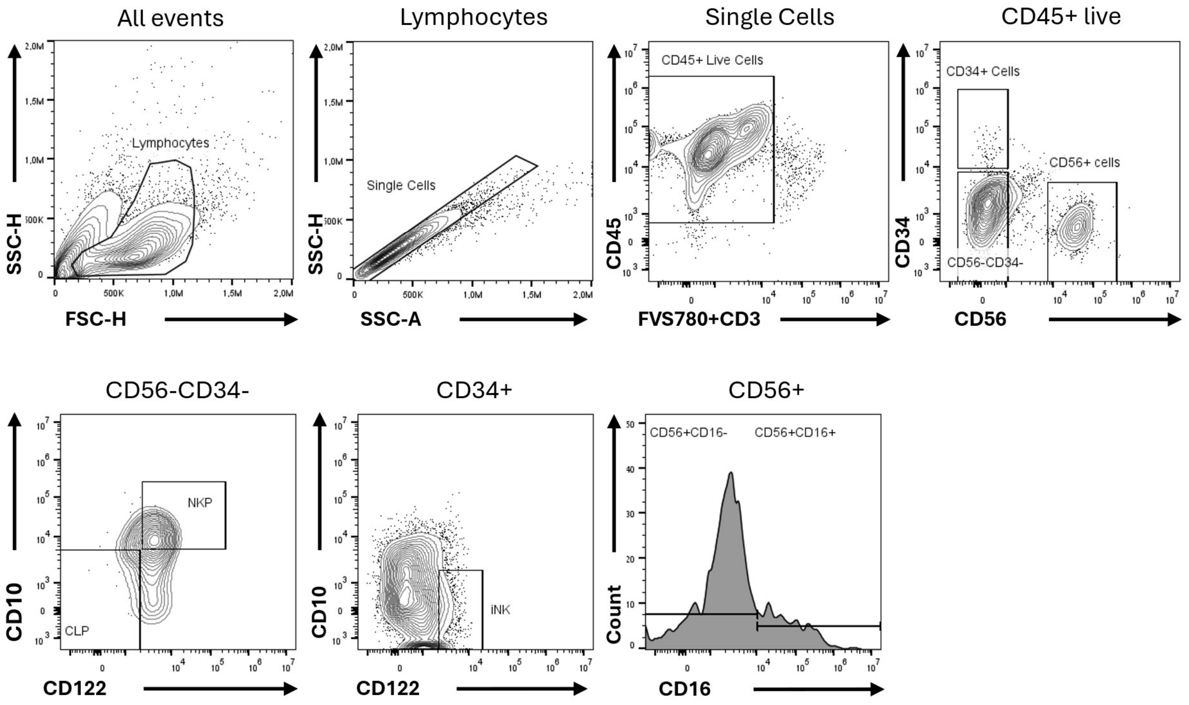


*Supplemental Figure 3. Flow cytometric analysis of progenitor populations during NK cell differentiation. NK cell panel successive gating strategy used for immunophenotypic identification of NK lineage progenitors. Plots are representative of cells analyzed on day 35 of NK cell differentiation.*
